# hyperTRIBER: a flexible R package for the analysis of differential RNA editing

**DOI:** 10.1101/2021.10.20.465108

**Authors:** Sarah Rennie, Daniel Heidar Magnusson, Robin Andersson

## Abstract

RNA editing by ADAR (adenosine deaminase acting on RNA) is gaining an increased interest in the field of post-transcriptional regulation. Fused to an RNA-binding protein (RBP) of interest, the catalytic activity of ADAR results in A-to-I RNA edits, whose identification will determine RBP-bound RNA transcripts. However, the computational tools available for their identification and differential RNA editing statistical analysis are limited or too specialised for general-purpose usage. Here we present `hyperTRIBER`, a flexible suite of tools, wrapped into a convenient R package, for the detection of differential RNA editing. `hyperTRIBER` is applicable to complex scenarios and experimental designs, and provides a robust statistical framework allowing for the control for coverage of reads at a given base, the total expression level and other co-variates. We demonstrate the capabilities of our approach on HyperTRIBE RNA-seq data for the detection of bound RNAs by the *N6*-methyladenosine (m^6^A) reader protein ECT2 in Arabidopsis roots. We show that `hyperTRIBER` finds edits with a high statistical power, even where editing proportions and RNA transcript expression levels are low, together demonstrating its usability and versatility for analysing differential RNA editing.

## 1 Introduction

RNA modifications constitute an important layer of regulation and diversification at the transcriptome and protein level [1, 2] in both animals and plants [3, 4, 5]. The study of RNA-editing, which refers to sequence changes made directly to RNA transcripts whilst the DNA sequence remains unchanged, has intensified in recent years along with the increasing availability of transcriptome-wide readout technologies such as RNA-seq [6, 7]. The most common type of RNA edit observed is adenosine to inosine (A-to-I), where I is sequenced as G. This type of RNA edit is mediated by the highly conserved adenosine deaminase enzyme ADAR and is thought to influence a variety of regulatory processes. In humans, ADAR1 is responsible for editing events at non-coding repetitive Alu regions [8] implicated in the innate immune response [9, 10]. ADAR2 editing occurs generally in protein coding regions, particularly in the brain, where their dis-regulation has been associated with neurological conditions [11, 12, 13]. Although the ADAR proteins are highly conserved over a wide taxa [14], they do not exist in plants, where C-to-U edits appear frequently in chloroplast and mitochondrial genes [15, 16].

RNA-editing is clearly observed in native contexts, but experimental technologies can also utilise editing by ADAR to introduce A-to-I edits, allowing for the identification of relevant transcripts. One example is the HyperTRIBE assay [17, 18] (targets of RNA binding proteins identified by editing), which can be used to find transcripts bound by RNA binding proteins (RBPs), by fusing a hyperactive variant of the catalytic domain of ADAR to a RBP of interest followed by measuring A-to-I editing levels in the vicinity of binding events. The HyperTRIBE assay can be used as an alternative or complement to purification-based methods like iCLIP (individual nucleotide resolution crosslinking and immunoprecipitation) [19], with the important difference that iCLIP in theory detects the exact locations of the binding site. However, due to biases inherent in CLIP based methodologies, including biases towards the most highly expressed genes [20], HyperTRIBE provides a powerful orthogonal methodology for transcript detection.

Analysis of these types of experiments are frequently based on direct comparisons between RNA-seq data sets, involving samples containing the RBP fused with ADAR and control-based samples without the fusion, and as such do not involve comparison with the corresponding sequenced DNA. Furthermore, there may also be batch or dosage based effects to be taken into account, for example large detected variation in ADAR abundance across HyperTRIBE samples could result in high variations in editing levels, making it difficult to detect statistically significant sites. Currently, the tools available for the statistical analysis of experiments such as HyperTRIBE and the subsequent annotation of the resulting regions remain limited or too specialised for general-purpose usage.

Here we present a set of flexible tools for the detection and annotation of RNA-editing sites between two conditions with available RNA-seq data (available at https://github.com/sarah-ku/hyperTRIBER). The `hyperTRIBER` R package leverages the DEXseq framework [21] to find sites whereby editing levels at a given nucleotide site is statistically different between the two conditions, whilst controlling for coverage of reads at the given base, the total expression level and other co-variates using a flexible generalised-linear model (`glm`) based framework. This approach allows for a robust, powerful and high-throughput statistical set up, which can effectively involve replicated samples. We demonstrate our approach on the application of HyperTRIBE for the detection of bound RNAs by the *N6*-methyladenosine (m^6^A) reader protein ECT2 in Arabidopsis roots [22], through identification of A-to-I edits introduced in the vicinity of the RNA binding location of ECT2. We show that our approach has high power to detect these A-to-I editing sites, even where the editing proportion is very low, as observed in the vast majority of cases. We further evaluate, through down-sampling, the sensitivity of `hyperTRIBER` to detect ECT2-bound targets and HyperTRIBE edit sites in low expressed transcript. Finally, we demonstrate that the flexible `glm` framework can be used to account for variation in ADAR levels across samples, resulting in a large increase in statistically significant sites, in the case of detecting ECT2- and ECT3-bound targets between two different genetic backgrounds [23].

Taken together, `hyperTRIBER` is a robust and versatile suite of tools for the detection of differential RNA editing at high sensitivity even at low editing proportions and low expressed RNA transcripts, which we expect will be a useful tool for the analysis of HyperTRIBE data in particular but also for studying native RNA editing events.

## 2 Methods

### 2.1 hyperTRIBER differential RNA editing work flow

Here we describe the work flow of `hyperTRIBER` (**Figure 1**) for differential RNA editing, including individual library processing, statistical comparisons and annotations of results. Further details and a **vignette of example analysis** can be found at the package GitHub repository (https://github.com/sarah-ku/hyperTRIBER).

**Figure 1:**
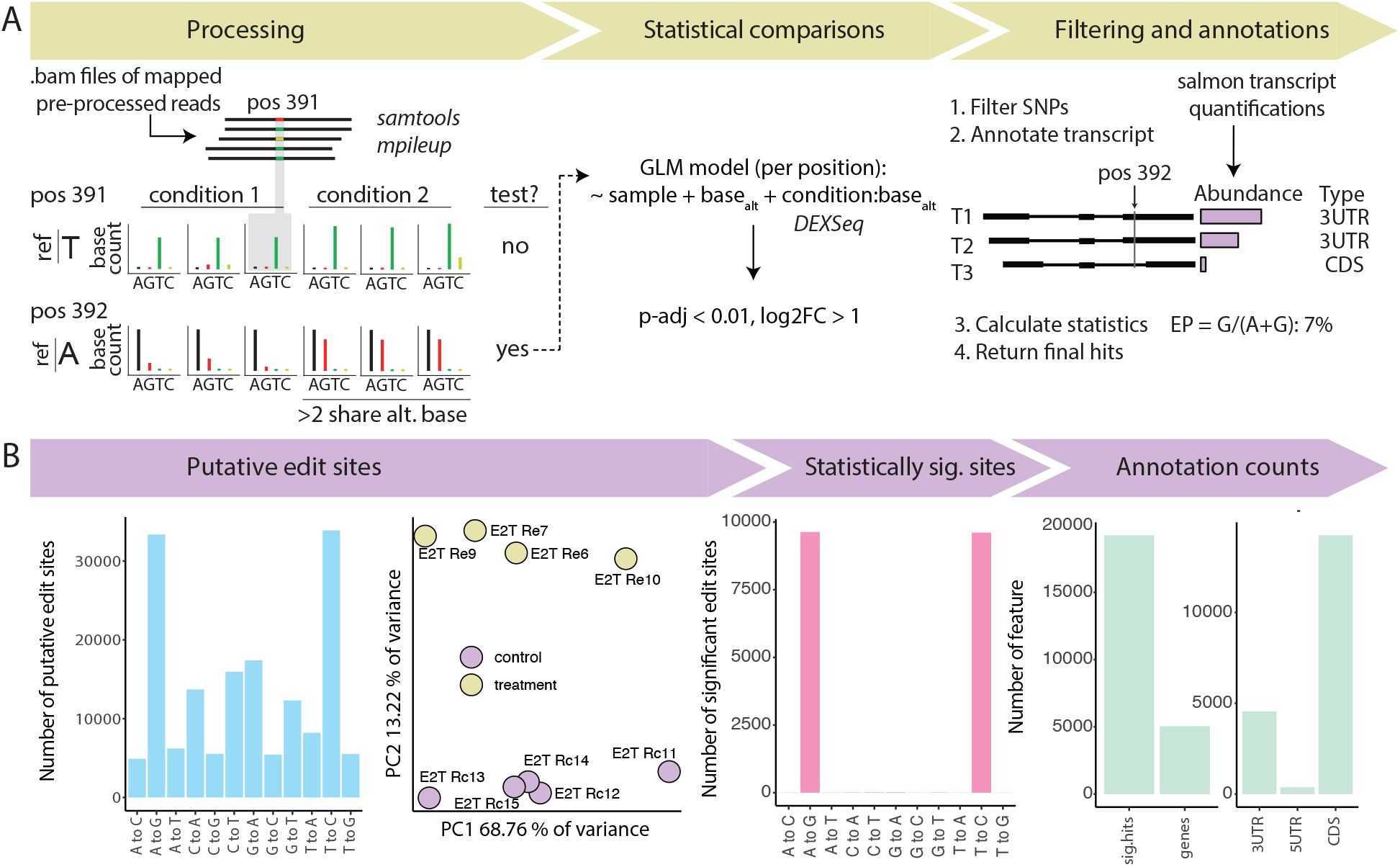
Overview and application of the three major steps of the hyperTRIBER analysis workflow for differential RNA editing analysis. **A**: Starting with processed and mapped .bam files, all positions with an alternative allele mapped in at least one sample are extracted and further filtered down to a list of potential edit sites by requiring replicate sharing. All potential edit sites are then tested for statistical significance using a GLM approach based on the same concept as differential exon usage (DEXSeq, [21]). Positions are then filtered and annotated using our custom strategy. **B**: An example application of the workflow to HyperTRIBE data for detecting ECT2 binding events in *Arabidopsis Thaliana* [22]. Left to right: distribution of putative editing sites according to editing type; principical component analysis of putative editing proportions; distribution of significant editing sites according to editing type; post-annotations counts according to the number of significant sites, the number of annotated genes and the number of significant sites annotating to the the 3’ UTR (3UTR), 5’ UTR (5UTR), or coding sequence (CDS).

#### 2.1.1 Step 1: individual library processing

The `hyperTRIBER` work flow starts with mapped BAM files of processed reads, one for each sample. These are simultaneously run through the function `mpileup` from `samtools` [24] and the output is then transformed (`extractPositionsFromMpileup.pl`, stranded and unstranded versions available) into human interpretable base counts for each position, whereby there is at least one read mapping to an alternative allele (that is, different from the reference genome supplied) in one sample. Note that for paired end stranded data, extra processing is required to arrange the BAM files into forward and reverse strands, instead of forward and reverse mate pairs (code provided), before pile-up data is produced. In order to run `mpileup`, a reference genome to compare the bases in the reads to is required, and although the final comparisons between the two conditions are technically reference free, it can be useful to check that the dominant base in at least one of the conditions is the same as the reference.

To complement the analysis, individual (gzipped) `fastq` files are also quantified using the tool `salmon` [25] for two purposes. First, quantification of potential modifiers such as ADAR over the samples can be used in the statistical test as presented in Step 2 below. Secondly, quantification averaged over samples in the control case (or according to the preference of the user) can be used in the gene annotations stage in order to prioritise possible transcripts within a gene and provide the most accurate possible annotations (see annotations description below).

#### 2.1.2 Step 2: statistical comparisons

For a given list of possible edit types of interest (e.g. A-to-G or C-to-T), the `hyperTRIBER` R function `restrictPositions` is used to select for possible sites potentially containing one or more of these edits, such that possible edits are shared across a given number of replicates, with a given number of reads covering the edit in each of the replicates.

In the statistical modelling step `hyperTRIBER` borrows functionality from the R package `DEXSeq` [21]. This package is designed for the concept of differential exon usage, whereby one can formally test if the proportions of reads mapping to a given exon are significantly different between two conditions, controlling for the overall number of reads mapping to the gene and any potential co-variate. In the current context, exons are instead replaced by bases (A, C, T, G) and the overall number of reads mapping to the gene is replaced by the overall number of reads mapping to the specific potential editing site (position). The relative usage of a base is thus defined as the fraction of reads mapping a specific base at a given position among the total number of reads mapping to any base at that position.

For the modelling, there currently is functionality in `hyperTRIBER` for two types of models, where the simplest model is:

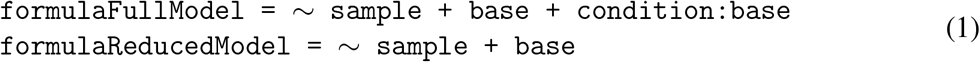

where `condition` refers to the two given conditions (for example, treatment and control in an experimental set-up), `sample` refers to an individual sample and `base` refers to the target base (for example G in the case of an A-to-G edit). In this way, the model (1) is testing whether the proportion of reads at the edit base is significantly higher in one condition than the other, controlling for the overall read coverage in the samples.

In the case where one would like to adjust for the levels of a specific modifier, one can use the following more complex version of the models:

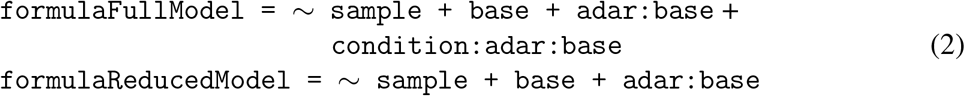

Here, `adar` is the given modifier protein and `adar:base + condition:adar:base` means that the editing proportion of the given base is affected by how much of the modifier protein there is in the system, which could also depend on the condition (interaction). The purpose of using this model (2) would be to reduce the variation in editing within each condition attributed to different levels of the modifier protein, thereby increasing the power to detect a significant difference.

The two models above (1 and 2) are run within the `DEXSeq` framework using a wrapper function supplied in the `hyperTRIBER` R package (instructions available on the GitHub repository). The annotation functions in the next step are used to interpret the output from `DEXSeq` in the context of RNA editing.

#### 2.1.3 Step 3: annotations

The output from the statistical comparisons in Step 2 is provided in a user-friendly `GRanges` object, holding information about whether each base has changed significantly in editing proportion. In the vast majority of cases, only a single edit type is considered, hence if there are two edits at the same site, the most significant edit is prioritised. The reference base is calculated as the dominant base at that position in the condition designated the control and the edit base is the significant base from the other condition. The editing proportion is calculated in each condition separately as Edit*/*(Ref + Edit), where Edit and Ref represent the counts of the edit and reference base respectively at the given position.

Annotations of edited sites are made in a manner prioritising transcript expression levels (**Figure 2**). First, the position of a site is overlapped with the relevant GTF file, and all transcripts overlapping the position are ordered according to transcripts per million (TPM) abundance (supplied as a vector). For each transcript, the position is assigned a feature of either 5UTR (5’ untranslated region), CDS (coding sequence) or 3UTR (3’ untranslated region), and its 5’-3’ proportion along the feature is calculated. All transcripts are given separated by commas, but in many downstream analyses it is often most relevant to use the most expressed transcript.

**Figure 2:**
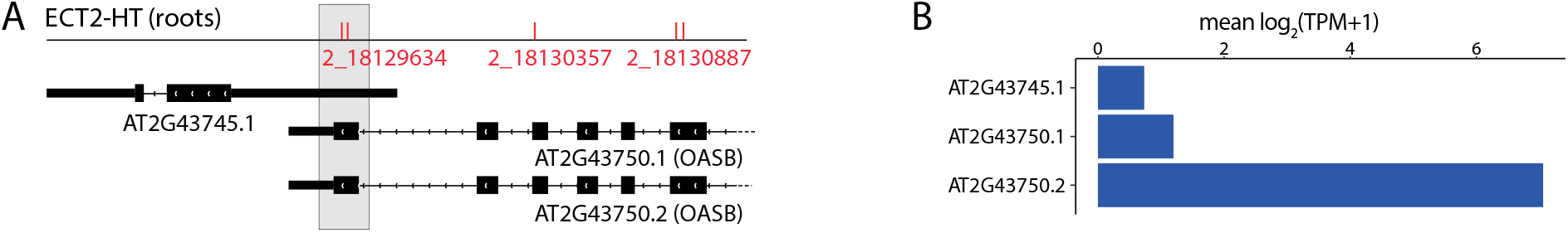
Demonstration of annotation strategy on significant HyperTRIBE editing sites overlapping multiple genes and transcripts. **A**: Significant A-to-G editing sites in ECT2-HT roots samples (red dashes) found using hyperTRIBER and located ambiguously within either AT2G43745 or OASB (leftmost sites in grey shaded area). **B**: Analysis of abundance shows the second transcript of OASB (AT2G43750.2) to be far the most expressed, supporting the annotation of these sites to this transcript.

### 2.2 Analysis of HyperTRIBE data

#### 2.2.1 Preprocessing of data

Paired-end reads of HyperTRIBE RNA-seq libraries [22, 23] (European Nucleotide Archive, PRJEB44359) were trimmed using Trimmomatic version 0.38 [26], before being mapped to the Arabidopsis Thaliana TAIR10 reference genome using STAR [27], whilst allowing for mismatches. Reads were further filtered based on quality using Picard tools (http://broadinstitute.github.io/picard/). The resulting SAM files were then converted to BAM files and sorted, before being run through the first set of the `hyperTRIBER` work flow described above.

#### 2.2.2 Analysis of HyperTRIBE data using the hyperTRIBER work flow

We filtered positions selected from the base counting stage further by requiring that at least 4 out of 5 replicates supported the potential edit. After running the model (model 1), we considered only significant positions (adjusted p-value ≤ 0.01) with a log_2_ fold change of at least 1, meaning that only positions which were edited significantly higher in the *ECT2-FLAG-ADAR* samples, which correspond to samples where the catalytic domain of *ADAR* is fused to *ECT2* and based on an *ECT2* knockout single mutant background *ect2-1*, were counted. Because we were only interested in positions associated with editing by ADAR, we selected only those which represented an A-to-G edit or a T-to-C edit (revealing hits on positive and negative strands, respectively; unstranded protocol). In order to further filter hits, we removed cases where the control samples, which include non-native ADAR but in absense of fusion (*FLAG-ADAR*), had a dominant base that did not agree with the reference genome base. We also filtered out cases where the editing proportion was very close to or exactly one, since close inspection of these hits showed that the likely cause was a single nucleotide variant (SNV) appearing in either of the sets of lines. Finally, genes were annotated according to Araport11 and transcript expression was based on the output from `salmon` from the *FLAG-ADAR* control samples. Positions were also assigned a number between 0 and 1 according to the proportion of the way through the given feature (5UTR, CDS or 3UTR) they were annotated to and used to generate a metaGene plot.

In order to compare effects of running models (1) and (2), we used HyperTRIBE datasets comparing fusion samples with a single mutation of the gene corresponding to the given protein of interest (*ect2-1* or *ect3-1* backgrounds for the respective RBPs ECT2 or ECT3) with matched fusion samples for the same protein of interest but based on a triple mutant knockout background of the genes *ECT2, ECT3*, and *ECT4* (*te234*) (see redundancy analysis [23]). Variation in editing levels by ADAR was calculated using the coefficient of variation (mean/standard deviation) across samples of detected transcripts mapping to *ADARclone* from the `salmon` output. Models comparing the single versus triple mutant set-up for each of ECT2 and ECT3 RBP separately were run using both models (1) and (2), and the numbers of significant hits and corresponding genes counted from the application of the annotation pipeline.

#### 2.2.3 Down-sampling and saturation analysis

In order to measure the sensitivity of our analysis approach to read depth, HyperTRIBE reads for ECT2 in roots were randomly sub-sampled three times to 5, 10, and 20 million reads. For each of the three down-sampled sets, the analysis was performed in an identical way as with the set with the full number of reads (∼ 40 million). The number of significant positions were counted for each of the sets, together with the total number of covered genes and the average number of hits per gene.

In order to calculate support from m^6^A modifications in identified transcripts, m^6^A site locations derived from Oxford nanopore data [28] were considered. The proportion of transcripts found in HyperTRIBE data that were supported by at least one m^6^A site per transcript was then calculated.

For the saturation analysis, genes found in the full (non-sampled) data were annotated according to whether they were also found in each of the three results sets based on the subsets of reads. A `glm` model was fit in `R` in order to determine whether the gene could be identified at a given gene expression level and editing proportion. Logistic regression curves were plotted as a function of gene expression, separately according to possible editing proportions and whether there was a single or multiple edit positions found in the gene.

## 3 Results

### 3.1 Application of the hyperTRIBER tool suite to HyperTRIBE data

We considered Arabidopsis HyperTRIBE RNA-seq data in roots [22] generated with the aim to identify transcripts bound by ECT2, a reader protein that detects and binds to m^6^A base modifications on transcripts, with potential implications to transcript stability and development [5, 29]. The control samples (*FLAG-ADAR*) were derived from transgenic plants that expressed artificially (or ectopically) introduced ADAR (note that ADAR is not native to Arabidopsis), such that ADAR is free to edit adenosines randomly. The controls were compared to samples where ECT2 was fused to the catalytic domain of ADAR (*ECT2-FLAG-ADAR* fusion samples, see methods or [22]) and thus editing should be more highly concentrated around locations where ECT2 binds to transcripts.

We ran the `hyperTRIBER` suite of tools on HyperTRIBE RNA-seq data from roots in Arabidopsis and found 19,242 positions after filtering for possible SNVs (described above) and positions that were not A-to-G (or T-to-C on the negative strand) (**Figure 3**). These hits were spread over a total of 5,053 unique genes (annotations based on Araport11), resulting in an average of 3.8 hits per gene, with a minimum of 1 and a maximum of 113 in the case of gene *PAB2*, (*AT4G34110*). Overall, editing proportions were very low, possibly explained by *ECT2* gene being expressed only in a limited sub-population of cells within roots [22]. Thus, we hypothesised that such cases would be hard to detect if the sample library depth was reduced.

**Figure 3:**
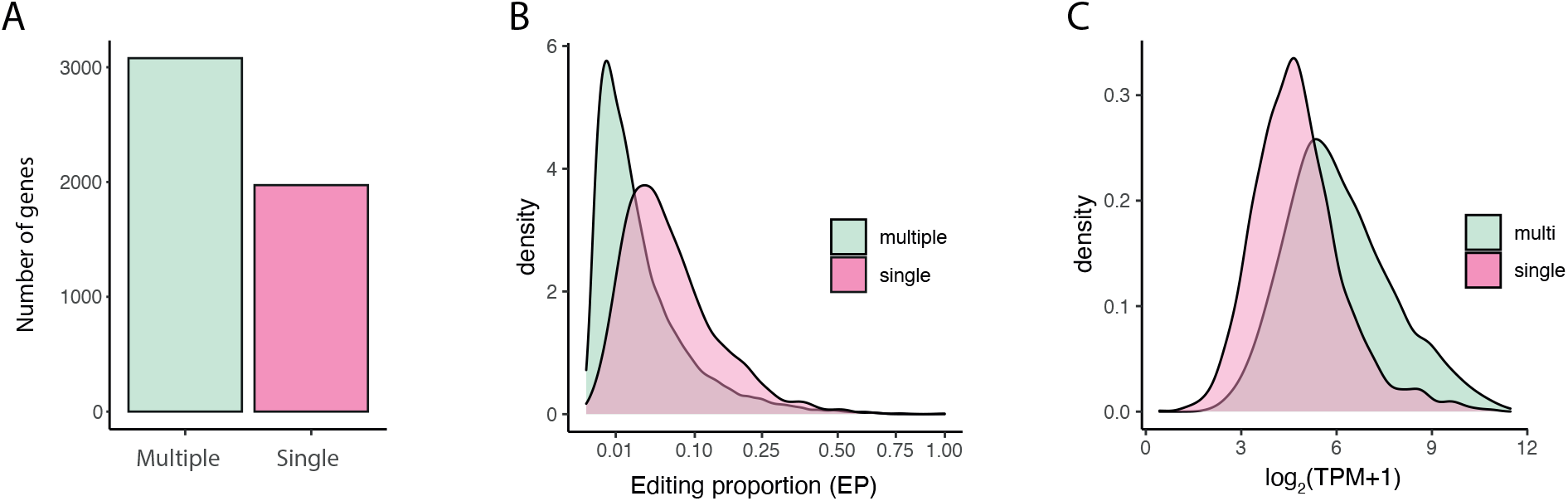
Single versus multiple edit site genes and their relationships with editing proportions and expression. **A**: Number of genes annotated to significant HyperTRIBE sites, according to whether multiple sites or a single site was found annotated to the gene. **B**: Density plot of editing proportions (EP) of significant HyperTRIBE sites, split according to whether they annotate to a single or multiple site gene. **C**: Density plot of transcript expression (log_2_(TPM+1)), according to whether the gene has multiple editing sites or a single editing site.

As expected, down-sampling the reads found fewer individual positions and genes (**Figure 4A,B**), but also fewer hits per gene (**Figure 4C**), suggesting that additional reads resulted in the pooling of positions often within the same gene. Support from m^6^A nanopore data [28] suggested that over 80% of transcripts detected using 5 million reads also had an m^6^A site predicted from nanopore reads (**Figure 4D**). This percentage decreased with an increasing number of reads, likely because genes detected using low sequencing depth are likely to be the most expressed, whose modifications are easier to detect using nanopore sequencing.

**Figure 4:**
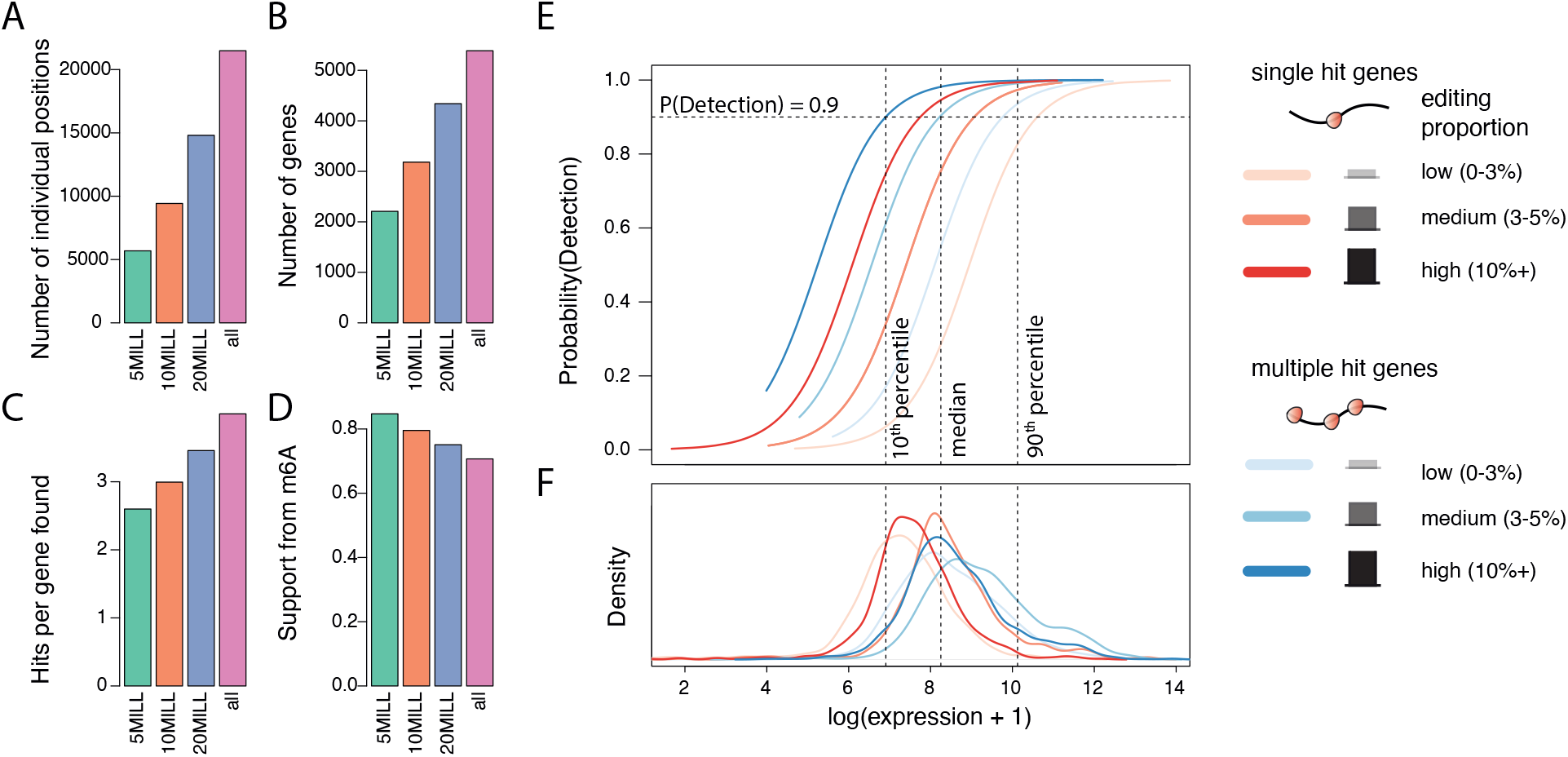
Subsampling analysis and the ability to detect positions with low editing proportions. **A-D**: Summary of positions found for Arabidopsis HyperTRIBE data using hyperTRIBER. Pink indicates the number of positions (A), genes (B), hits (C) and m^6^A support proportion (D) for all reads. The other colours represent the same statistics, after down-sampling the full set of reads to 5, 10 or 20 million reads. **E-F**: The horizontal axis shows log(expression+1), with the dashed lines representing the 10^th^, 50^th^ and 90^th^ percentiles of the expression of all genes in the set. The top panel (E) represents the probability of detecting a gene containing at least one ADAR-edited site, further split according to editing proportion (low, medium, high) and whether the gene contained a single hit or multiple hits. The bottom panel (F) displays their densities.

It is also relevant to investigate the sensitivity of the analysis according to expression levels together with varying editing proportions (**Figure 4E,F**). For example, the probability of detecting at least one edit site in a multi-hit gene with an editing proportion of at least 10% is 0.9 for a gene in the 10^*th*^ percentile of expression, and close to 1 for a gene over median expression. Conversely, genes with a low editing proportion (0-3%) had much lower detection rates, particularly in the cases of lowly expressed genes. Thus, it is likely that HyperTRIBE has missed some very lowly expressed, likely cell type specific genes, which could well be targets of the reader protein ECT2.

Finally, we tested the effect of adjusting for levels of the editing enzyme ADAR (model 2) on the detection of differentially edited positions in HyperTRIBE data for detecting the binding of the RBPs ECT2 and ECT3 to m^6^A sites [22, 23]. For each protein, the scenario where the respective *ECT2* or *ECT3* gene was knocked-out (single mutant) was compared to the scenario where all three of *ECT2, ECT3*, and *ECT4* were knocked-out (triple mutant) (see methods). Positions significantly edited in the triple mutant setting versus the single mutant setting were indicative of redundant sites, which would not normally be bound by the tested protein in the native context. In general, the levels of detected ADAR varied more in the ECT3 samples, as measured by the coefficient of variation, as well as in the single mutant for each experiment (**Figure 5A**). When applying the adjustment model (model 2), greater numbers of significant differential editing was found compared to the standard model (model 1) (**Figure 5B**). This effect was particularly pronounced in the ECT3 data, probably reflecting the comparatively high ADAR covariance amongst biological replicates (**Figure 5A**). Furthermore, the extra positions also translated into more affected genes, for both ECT2 and ECT3 datasets (**Figure 5C**).

**Figure 5:**
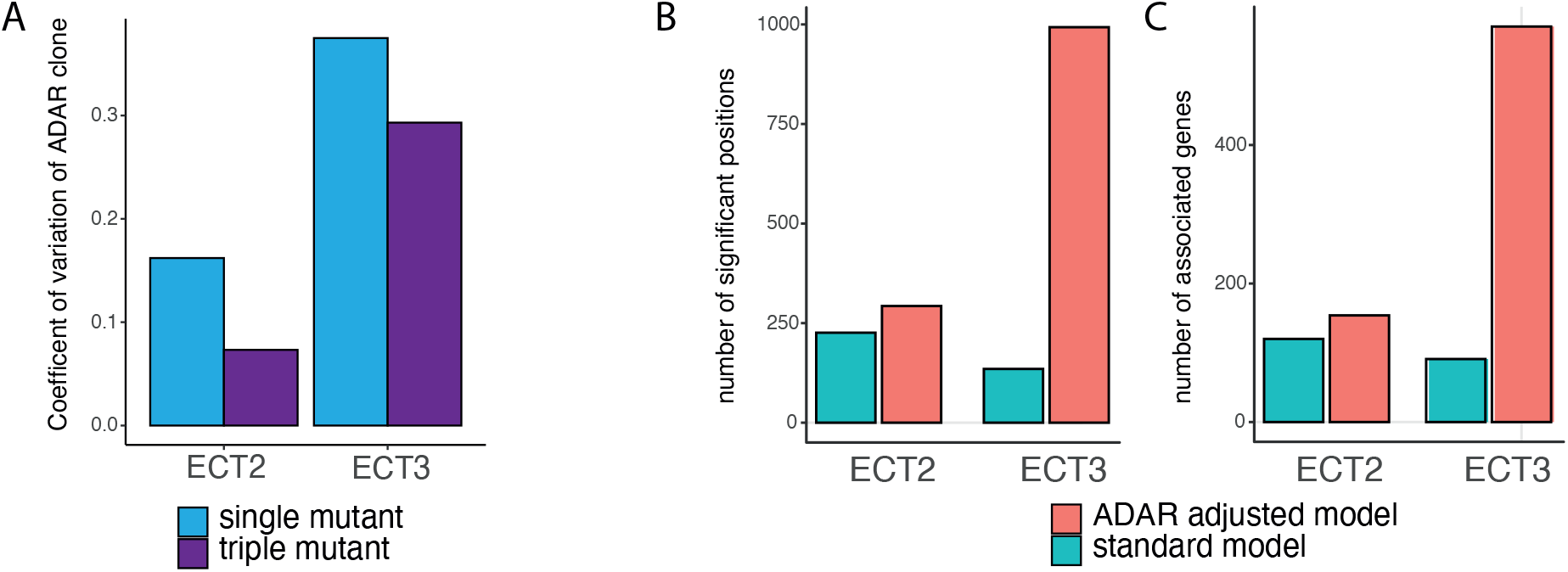
Application of ADAR adjusted model for comparing single and triple mutant HyperTRIBE samples for each of ECT2 and ECT3 in *Arabidopsis roots*. **A**: Coefficient of variation across samples, within the four experiments. Note larger coefficient of variation in ECT3 samples. **B**: Number of significantly differentially edited positions between single and triple mutant experiments for each of ECT2 and ECT3 roots datasets. **C**: Number of associated genes, based on results from using hyperTRIBER. Two models were considered in the analysis, as described in Methods. The standard model (model 1) adjusts for expression levels at the given position and tests for significant differences in editing base proportion. The ADAR adjusted model (model 2) controls for extra variation in editing levels due to differences in ADAR abundance across samples.

## 4 Discussion

With increasing quantities of RNA-seq data, RNA-editing is becoming a more widely studied field of research, but tools to detect and effectively analyse editing events are still scarce or ad hoc. Here we have presented a framework, `hyperTRIBER` for detecting differential editing in RNA between pairs of conditions. Our framework is built upon the `DEXSeq` framework which is fast, efficient, powerful, and makes good use of replicates. It is also flexible, which allows for the adjustment of potential modifiers, such as ADAR, to further increase the power of the test. We also provide a framework for annotating the positions, as well as further functions for downstream analysis, which we continue to develop.

We demonstrated our framework on HyperTRIBE datasets from our study involving the detection of transcripts bound by the m^6^A reader protein ECT2 or ECT3 in *Arabidopsis* roots [22, 23]. We note that whilst the framework was originally designed for this type of data, and named as such, we envisage it’s usefulness in detecting native RNA editing events. Here we have shown that `hyperTRIBER` has power to detect statistically significant sites and thus transcripts, despite inherently low editing proportions in the HyperTRIBE data. We also demonstrate `hyperTRIBER`’s ability to detect positions at various sampling depths and expressions, arguing that whilst we are able to detect lowly expressed transcripts which were otherwise missed using iCLIP [22], there may still be potential to find further, rare, transcripts bound by the RBP by using greater sequencing depths. Applying extended models controlling for variations in ADAR was also shown to be a powerful approach, finding more significant sites and corresponding transcripts bound by the RBP than the standard model. We note, however, that the ADAR adjusted model required more computational resources than the standard model.

We further note that the `hyperTRIBER` package does not account for known SNVs, which could show up between individuals in a population. This is particularly relevant to consider when studying native RNA editing. However, we also observed a few potential false positive cases in the analysed HyperTRIBE data arising between the wild-type ADAR lines and the ADAR fusion lines, which began as genetically identical. These tended though to have editing proportions close or equal to one, and were often distal from other editing sites, raising suspicion about their credibility, and thus were straightforward to filter out. It is therefore important to evaluate among `hyperTRIBER` results which events are likely to be true editing events, and which are from other causes. Since editing sites often appear to cluster in similar regions, a potential filtering method could therefore be to require that there are multiple sites in a given transcript or interval, and filter out the rest (as used for example in [13]).

In conclusion, we present `hyperTRIBER`, an R package that offers a flexible and powerful framework for detection of RNA edit sites, which we believe will be useful for both analysing HyperTRIBE data and native RNA editing events.

## 5 Acknowledgements

We thank Laura Arribas-Hernández, Carlotta Porcelli, and Peter Brodersen for fruitful discussions during the development of `hyperTRIBER`, and Laura Arribas-Hernández for providing feedback on the manuscript.

## 6 Funding

This work was supported by the European Research Council (ERC) under the European Union’s Horizon 2020 research and innovation programme [grant number 638173].

## Notes

https://github.com/sarah-ku/hyperTRIBER

### Competing Interest Statement

The authors have declared no competing interest.

